# *Wolbachia*-mediated parthenogenesis induction in the aphid hyperparasitoid *Alloxysta brevis* (Hymenoptera: Figitidae: Charipinae)

**DOI:** 10.1101/2025.06.30.662338

**Authors:** Jonathan Dregni, Amelia R. I. Lindsey, Mar Ferrer-Suay, Sabrina L. Celis, George E. Heimpel

**Author notes:** Address correspondence to George Heimpel. **Author Contributions**: J.D.: conceptualization, antibiotic experiment; insect and plant rearing, writing; A.R.I.L.: molecular analyses, writing, funding; M. F.-S.: hyperparasitoid identification, worldwide sex ratio data, writing, funding; S.L.C.: hyperparasitoid rearing, Fig. 1, writing, G.E.H. conceptualization, writing, funding. Authorship order: First author: experimental work and planning; last author: principal investigator.

## Abstract

Thelytokous parthenogenesis (thelytoky), in which females can produce female offspring without mating, can be caused by parthenogenesis-inducing endosymbiotic bacteria in the genus *Wolbachia*. This interaction is well known in hymenopteran parasitoids, where unfertilized eggs typically develop as males via haplo-diploidy in the absence of parthenogenesis-inducing bacteria. We report on a case of thelytoky in *Alloxysta brevis* (Thomson) (Hymenoptera: Figitidae), a globally widespread aphid hyperparasitoid. A previous study had shown that sex ratios of this species collected in Minnesota (USA) were extremely female biased, and we found here that unmated females reared from field-collected hosts produced female offspring without exposure to males. This result demonstrated thelytoky, and we tested for the role of bacterial endosymbionts by comparing offspring production of unmated females fed the antibiotic rifampicin to offspring production of control females not fed antibiotics. Antibiotic-fed females produced almost exclusively male offspring, and control females produced mainly females. This result showed that antibiotic treatment facilitated male production by unmated *Alloxysta brevis* females, thus implicating bacterial symbiosis in the expression of thelytoky. We then used molecular analyses to determine the identity of the symbiont. These analyses identified a *Wolbachia* strain from supergroup B, and excluded other bacteria known to mediate parthenogenesis induction, such as *Cardinium* and *Rickettsia*. While *Wolbachia* had been previously detected by molecular analysis in this species, these are the first experiments demonstrating *Wolbachia*-mediated parthenogenesis in the figitid subfamily Charipinae.

**IMPORTANCE:** Parthenogenesis induction in insects can have important environmental and economic consequences. This is especially true if pests or their natural enemies are affected. The case of *Alloxysta brevis* is of particular interest, as this species is a hyperparasitoid of aphids, meaning that they attack and kill primary parasitoids of aphids. The populations of many species of pest aphids are controlled by primary parasitoid species, and hyperparasitoids thus have the potential to interfere with this mechanism of control. The role of hyperparasitoid parthenogenesis in the suppression of aphids by primary parasitoids remains unexplored. Thus, the results of this set of studies provides a starting point for determining whether parthenogenesis-inducing *Wolbachia* in hyperparasitoids should be expected to improve or hinder biological control of pest aphids by primary parasitoids. The focus on *Alloxysta brevis* as a model for these questions could be particularly instructive, since it is a species of worldwide distribution that is involved in numerous economically important aphid-parasitoid interactions.

## INTRODUCTION

Many species of parasitoid wasps exhibit thelytokous parthenogenesis (i.e., thelytoky), in which unmated females produce viable female offspring. Thelytoky can arise either from genetic mechanisms or from the actions of parthenogenesis-inducing endosymbiotic bacteria (1–3). In the latter case, the presence of bacteria can alter meiosis or mitosis to cause the production of diploid embryos in the absence of fertilization, instead of haploid embryos that would develop as males under haplo-diploid sex determination (4, 5). In parasitoid wasps, symbiont-induced parthenogenesis has been documented in the hymenopteran superfamilies Chalcidoidea, Cynipoidea, Ichneumonoidea and Platygastroidea (2, 6). While most documented cases of symbiont-induced parthenogenesis can be attributed to bacteria in the genus *Wolbachia*, other bacteria such as *Cardinium* and *Rickettsia* have also evolved parthenogenesis-inducing mechanisms (2).

Here, we determine whether the aphid hyperparasitoid *Alloxysta brevis* (Thomson, 1862) (Hymenoptera: Cynipoidea: Figitidae: Charipinae) exhibits symbiont-induced thelytoky, and if so, which microbe(s) are involved. Hyperparasitoids are parasitoids that attack other parasitoid species (7), and *Alloxysta brevis* is a broadly distributed species that uses a broad range of aphid parasitoids as hosts (8–11). We encountered *Alloxysta brevis* attacking soybean aphids, *Aphis glycines* Matsumura (Hemiptera: Aphididae) already parasitized by the parasitoid wasp *Aphelinus certus* Yasnosh (Hymenoptera: Aphelinidae) in soybean fields of Minnesota, USA (12). Of 264 adult *Alloxysta brevis* individuals reared, 263 were females and one was a male (12). Such an extremely biased sex ratio suggests thelytoky, and we used an experiment comparing the offspring of antibiotic-fed females with offspring of control females to evaluate the hypotheses that thelytoky was, in fact, operating in *Alloxysta brevis*, and if so whether it could be attributed to endosymbiotic bacteria. Having found this to be the case, we used molecular analyses to identify the microbial taxon (or taxa) involved. The presence of *Wolbachia* DNA was previously reported from one *Alloxysta brevis* specimen from Norway (13). In this study we confirm the presence of *Wolbachia* and show that a symbiont is mediating thelytoky in this species.

## MATERIALS AND METHODS

### Natural history of *Alloxysta* hyperparasitism

*Alloxysta* species are obligate aphid hyperparasitoids that oviposit into developing larvae of primary aphid parasitoids, including members of the genus *Aphelinus* (Hymenoptera: Aphelinidae) and of the subfamily Aphidiinae (Hymenoptera: Braconidae) (7; see Fig. 1A). A single egg is likely laid in most cases, but even in the case of multiple eggs being laid by a single female, or superparasitism (when different females oviposit into the same host), only a single adult *Alloxysta* emerges. *Alloxysta* eggs remains dormant until the primary parasitoid has pupated within a cocoon formed by the exoskeleton of the dead aphid (the ‘mummy’). The *Alloxysta* larva then consumes the pupa of the primary parasitoid internally and emerges from the aphid mummy (7).

**Figure 1.**
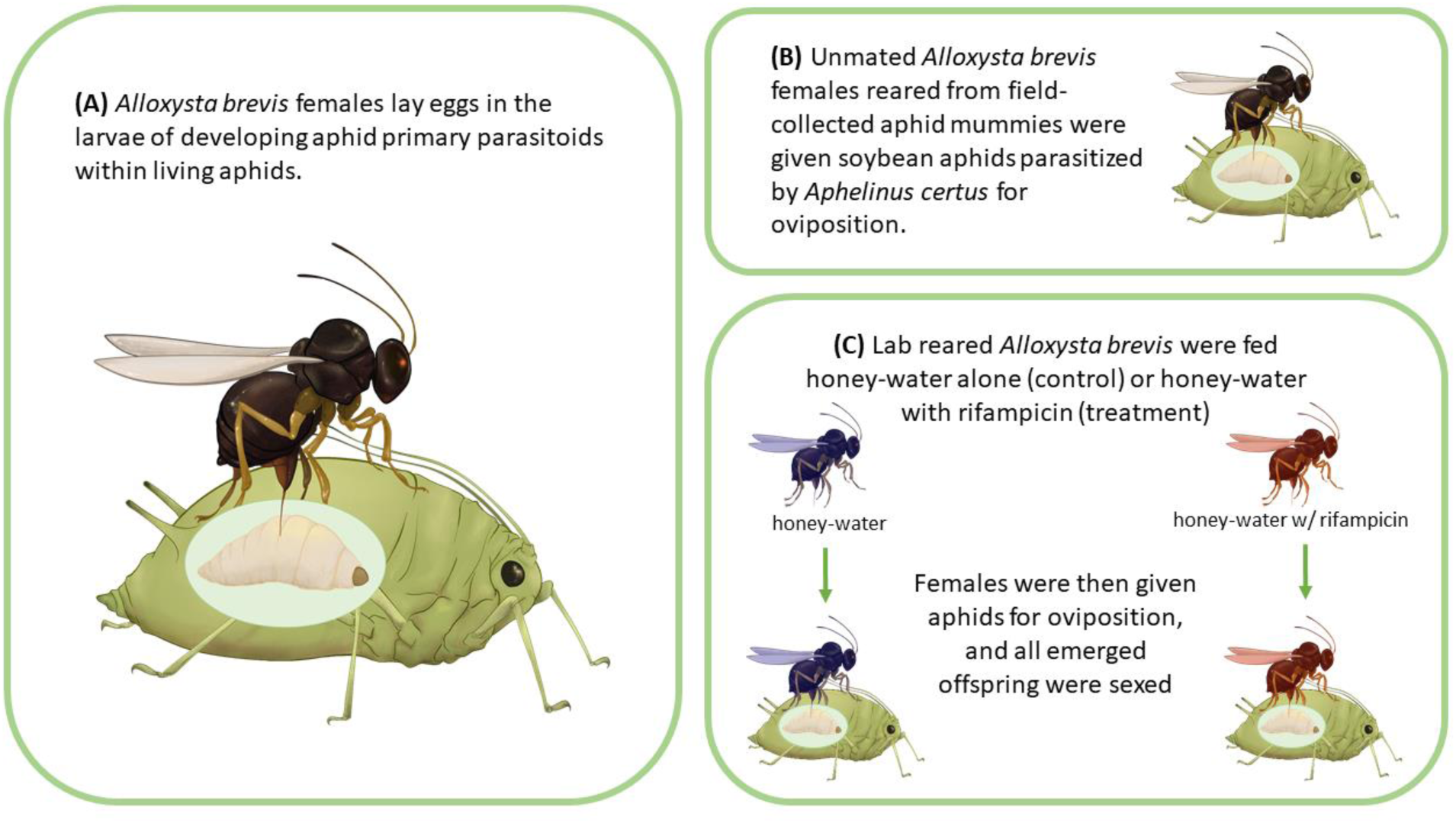
Schematics of *Alloxysta* oviposition and experimental design. **(A)** *Alloxysta brevis* ovipositing into an *Aphelinus certus* larvae within a soybean aphid. **(B)** Unmated *Alloxysta brevis* female exposed to soybean aphid containing an *Aphelinus certus* larva. **(C)** Schematic of protocol for antibiotic-feeding experiment. All figures drawn by Sabrina Celis.

### Review and analysis of *Alloxysta brevis* sex ratios

*Alloxysta brevis* is a globally distributed species, with samples collected in Africa, throughout Eurasia, as well as parts of the neotropics and North America. We compiled collections of this species used for taxonomic studies and tabulated the adult sex ratios associated with each collection. We report the proportion of females per geographical location and summed over regions. For any data sample consisting of greater than 20 individuals, we used a two-tailed binomial test to determine whether the sex ratio was significantly biased toward females or males.

### Field sampling and rearing hyperparasitoids

*Alloxysta brevis* were reared from field-collected *Aphelinus certus* mummies, collected along a 600 km transect within the soybean-growing area of Minnesota, USA, during summer 2018 using methodologies described by Casiraghi et al. (12). The mummies were then held in a growth chamber at 24.5°C +/- 1°C, at a 16:8 L:D photoperiod and observed daily for emergence. Hyperparasitoids emerging from *Aphelinus certus* mummies were identified to species by M. F.- S. and sexed by use of morphological characters using a key to the Charipinae (14). A colony was established by adding *Alloxysta brevis* females to plexiglass cages (30 x 38 x 36 cm) in which potted soybean plants (stages V1 – V3) were seeded with soybean aphids and *Aphelinus certus* (15).

### Assessment of thelytoky

To conduct an initial assessment of parthenogenesis in *Alloxysta brevis*, a sample of 23 females that emerged from field-collected *Aphelinus certus* mummies were held individually upon emergence and exposed to soybean aphids parasitized by *Aphelinus certus* from a laboratory colony. We infested potted soybean plants with soybean aphids and added 8 *Aphelinus certus* adults. After 6 to 9 days, we harvested leaflets containing at least 2 *Aphelinus* mummies and 20 aphids, an unknown number of which contained *Aphelinus certus* larvae of various ages. These aphids were exposed to *Alloxysta brevis* females by placing the leaflet petiolule into a centimeter of damp sand in a 30mL capped vial with a few droplets of 50% honey solution as a food source. We then added a single unmated *Alloxysta brevis* female to each vial and moved her to vials containing fresh leaflets with hosts as described above every two days (Fig. 1B). All aphid mummies were collected once they formed and placed singly into 0.6 ml microcentrifuge tubes, and observed for parasitoid emergence. Since none of the *Alloxysta brevis* females used in this study had had any exposure to males, the production of female offspring would be proof of thelytokous parthenogenesis (2, 4).

### Antibiotic treatment

We compared the reproduction of *Alloxysta brevis* females fed the antibiotic rifampicin to control females (Fig. 1C). Rifampicin was diluted in aqueous dimethyl sulfoxide at a concentration of 10mg/mL and then mixed with honey and water to a final concentration of 1mg/mL in 50% honey. Tiny (< 0.5 mm diameter) droplets of control (no rifampicin) or rifampicin honey were placed in 0.6 mL microcentrifuge tubes containing *Alloxysta brevis* females that had emerged within the previous 24 hours, and then held for 72 hours at 26.2°C +/- 0.5°C. After this period the *Alloxysta brevis* females were placed in soybean-leaf arenas with up to 20 *Aphelinus certus*-parasitized soybean aphids as described above. No antibiotic treatment was applied after the initial 72-hour feeding period, but all arenas were provided 50% control honey from this point onwards. *Alloxysta brevis* females were moved to fresh soybean leaflets containing parasitized soybean aphids as described above every 2 days for 12 days, resulting in 6 sequential assessments of reproduction for each experimental female. We evaluated 61 *Alloxysta brevis* females, alternately assigned either rifampicin honey (‘treated’), or untreated honey (‘control’), of which 19 treated and 16 control females reproduced during the course of the assay.

Mummies that developed on the leaves were isolated as described above and offspring species and sex were recorded upon emergence. We used two-tailed Fisher’s exact tests to determine whether there was a significant effect of the antibiotic treatment on male production, and whether there was a significant change in male production over the six time periods in the control treatment.

### Identification of bacterial endosymbiont

DNA was extracted from individual *Alloxysta brevis* specimens using HotSHOT (17) in a final total volume of 24 μL, as implemented previously for other microhymenopterans (18). *Wolbachia*-specific 16S PCRs were performed with WspecF and WspecR primers (19), and general bacterial full-length 16S PCRs leveraged 27F and 1492R primers (20) (Table 1). All PCRs were prepared in a final volume of 20 μL with 1 μL of DNA template, Q5® Hot Start High-Fidelity 2X Master Mix (New England BioLabs), and 500 nM of each primer, alongside positive and negative controls. Negative controls included no-template PCR reactions, and a negative control for environmental contamination consisting of the extraction buffer mix alone into which the wasp-handling paintbrush was dipped. Thermalcycling was conducted on a Mastercycler® nexus PCR cycler (Eppendorf®) with an initial denaturation of 2 minutes at 98°C, 35 cycles of amplification (see Table 1), and final extension of 2 minutes at 72°C. PCR products were separated with electrophoresis on a 1% agarose gel and visualized under ultraviolet light after staining with GelRed™ (Biotium) diluted to 3X in water. Prior to sequencing, PCR-products were cleaned using the Zymo DNA Clean & Concentrator® – 5 Kit (Zymo Research), and quantification was performed with a Qubit 4 Fluorometer and the Qubit 1X dsDNA HS Assay Kit (Invitrogen). Full-length 16S PCR products derived from eight individual *Alloxysta brevis* individuals were sequenced via the ‘Premium PCR Sequencing’ service by Plasmidsaurus using Oxford Nanopore Technology, alongside extraction negative controls. Fastq files generated by Plasmidsaurus were then analyzed following our own custom pipeline. Specifically, reads were first taxonomically classified using the Oxford Nanopore Technologies EPI2ME ‘wf-16s’ v1.1.3 workflow employing Kraken2 v2.1.2 (20). Then, reads corresponding to taxa not present in the negative extraction control were extracted with custom bash scripts and assembled with the EPI2ME ‘wf-amplicon’ v1.0.4 workflow which aligns and polishes reads to generate a consensus sequence using miniasm v0.3-r179 (21) and medaka v1.11.1. Consensus sequences and mapped reads (e.g., .bam files) were manually inspected in JBrowse (22) to check for chimeric assemblies. The final *Wolbachia* 16S sequence from *Alloxysta brevis* was queried against the NCBI GenBank database with blastn and default parameters. 16S sequences from top GenBank matches, plus those from a range of other well described *Wolbachia* strains (Table 2) were aligned with the secondary structure-aware SINA v1.2.12 aligner (23). Phylogenetic reconstruction was performed with IQ-TREE v1.6.11 (24) including model optimization and selection, outgroup specification, and 1000 rapid bootstraps.

**Table 1.**
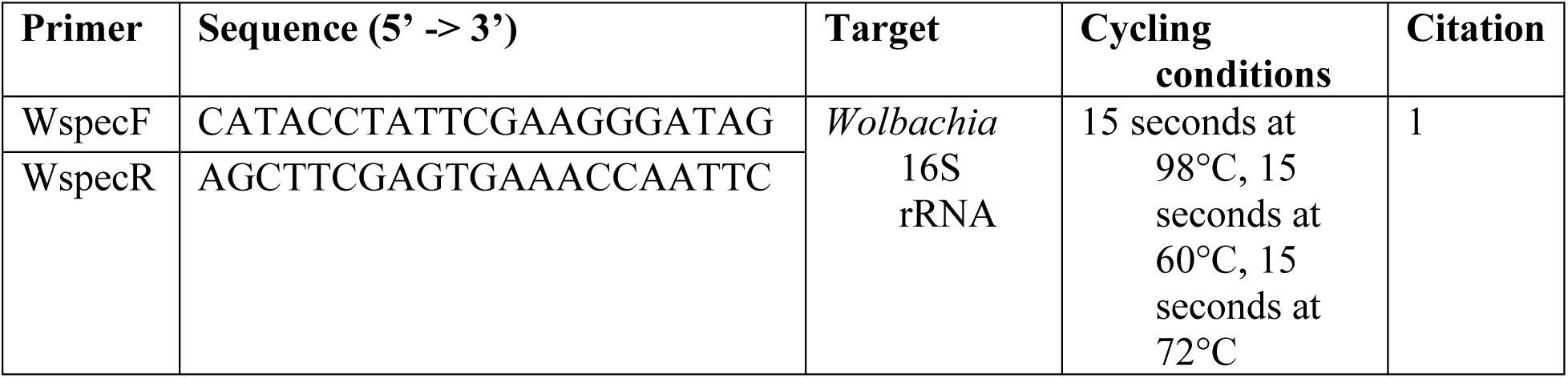
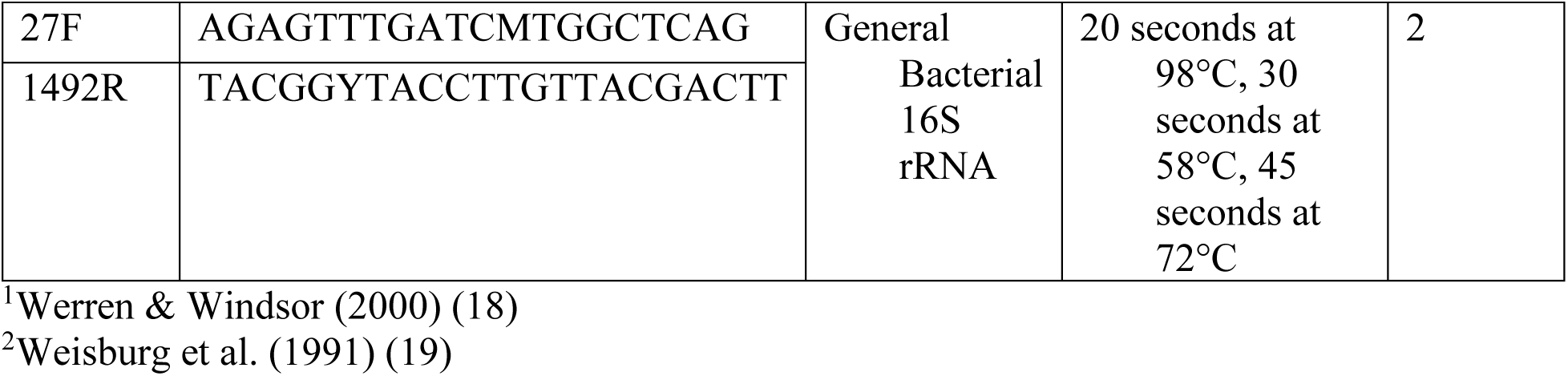
Information on primer pairs used to amplify *Wolbachia* sequences in *Alloxysta brevis*.

**Table 2.** 16S DNA sequences used in phylogenetic reconstruction of *Wolbachia* sequences associated with various host species indicating the *Wolbachia* supergroup (44, 45) and NCBI accession number.

| Strain | Host | Supergroup | Accession |
| --- | --- | --- | --- |
| wAlbA | *Aedes albopictus* | A | CP101657.1 |
| wAlbB | *Aedes albopictus* | B | CP041923.1 |
| wAu | *Drosophila simulans* | A | CP069055.1 |
| wBm* | *Brugia malayi* | D | AJ010275.1 |
| wBpra | *Bryobia praetiosa* | B | EU499317.1 |
| wCpunc | *Cyclophora punctaria* | B | OZ034749.1 |
| wDacB | *Dactylopius coccus* | B | LSYY01000126.1 |
| wDi | *Diaphorina citri* | B | CP048820.1 |
| wDmed | *Dolichovespula media* | B | CAMXBR010000003.1 |
| wHa | *Drosophila simulans* | A | CP003884.1 |
| wLcla | *Leptopilina clavipes* | B | QJHA01000026.1 |
| wMa | *Drosophila simulans* | B | CP069054.1 |
| wMau | *Drosophila mauritiana* | A | CP034335.1 |
| wMel | *Drosophila melanogaster* | A | CP046925.1 |
| wNo | *Drosophila simulans* | B | CP003883.1 |
| wOo* | *Onchocerca ochengi* | C | AJ010276.1 |
| wPip | *Culex quinquefasciatus* | B | AM999887.1 |
| wSuz | *Drosophila suzukii* | A | CAOU02000038.1 |
| wTei | *Drosophila simulans* | A | CP069052.1 |
| wTpre | *Trichogramma pretiosum* | B | LKEQ01000006.1 |
| wUni | *Muscidifurax uniraptor* | A | L02882.1 |
| CIXPIL1 | *Cixidia pilatoi* | B | OQ102143.1 |
| MALBOS1 | *Malenia bosnica* | B | OQ102152.1 |
| PARIOC1 | *Paracorethrura iocnemis* | B | OQ102158.1 |
| PENROR1 | *Pentastira rorida* | B | OQ102146.1 |

## RESULTS

### Global patterns of *Alloxysta brevis* sex ratios

The overall sex ratio of *Alloxysta brevis* adults collected across the globe was significantly female-biased (Table 3). This included cases of all-female samples of greater than 20 individuals in Africa, France and Russia, and cases of significant, extreme female-bias (> 90%) in Japan, Germany, the Nearctic region as a whole, and the state of Minnesota within the United States, as well as more modest (but still significant) female-biased samples from the Balkans (Table 3).

**Table 3.** Numbers of female and male *Alloxysta brevis* collected globally, along with the proportion female from each sample. Significant differences from equal sex ratios are indicated for samples comprising at least 20 individuals using two-tailed binomial tests: \**P* < 0.05, \*\**P* < 0.01, \*\*\**P* < 0.001. (9, 10, 12, 46–55)

| REGION/COUNTRY | FEMALE | MALE | PROP.<br>FEMALE | REFERENCE |
| --- | --- | --- | --- | --- |
| **AFRICA (sum)** | **28** | **0** | **1.00\*\*\*** | Ferrer-Suay et al. 2013d, 2018 |
| Morocco | 4 | 0 | 1.00 | Ferrer-Suay et al. 2018 |
| **ASIA (sum)** | **77** | **14** | **0.84\*\*\*** |  |
| China | 2 | 0 | 1.00 | Ferrer-Suay et al. 2013e |
| India | 3 | 2 | 0.60 | Ferrer-Suay et al. 2013e |
| Iran | 15 | 7 | 0.68 | Ferrer-Suay et al. 2013b, 2018 |
| Japan | 56 | 5 | 0.92*** | Ferrer-Suay et al. 2013e, 2018 |
| Thailand | 1 | 0 | 1.00 | Ferrer-Suay et al. 2013e |
| **EUROPE (sum)** | **542** | **110** | **0.83\*\*\*** |  |
| Balkan Peninsula | 153 | 85 | 0.64*** | Ferrer-Suay et al. 2013c; 2025 |
| Croatia | 1 | 0 | 1.00 | Ferrer-Suay et al. 2018 |
| Czech Republic | 11 | 1 | 0.92 | Ferrer-Suay et al. 2018 |
| France | 23 | 0 | 1.00*** | Ferrer-Suay et al. 2015, 2018 |
| Germany | 269 | 10 | 0.96*** | Ferrer-Suay et al. 2018; M. F.-S., unpubl. |
| Greece | 3 | 0 | 1.00 | Ferrer-Suay et al. 2018 |
| Italy | 13 | 7 | 0.65 | Ferrer-Suay et al. 2014b |
| Poland | 11 | 0 | 1.00 | Ferrer-Suay et al. 2020 |
| Russia | 23 | 0 | 1.00*** | Ferrer-Suay et al. 2018 |
| Serbia | 2 | 0 | 1.00 | Ferrer-Suay et al. 2018 |
| Spain | 3 | 1 | 0.75 | Ferrer-Suay et al. 2018 |
| Slovakia | 1 | 0 | 1.00 | Ferrer-Suay et al. 2018 |
| Slovenia | 9 | 4 | 0.69 | Ferrer-Suay et al. 2018 |
| Sweden | 4 | 0 | 1.00 | Ferrer-Suay et al. 2018 |
| Switzerland | 16 | 2 | 0.89 | Ferrer-Suay et al. 2018 |
| **NEOTROPICAL (sum)** | **8** | **1** | **0.89** |  |
| Ecuador | 1 | 0 | 1.00 | Ferrer-Suay et al. 2013d |
| Guatemala | 1 | 0 | 1.00 | Ferrer-Suay et al. 2013d |
| Mexico | 6 | 1 | 0.86 | Ferrer-Suay et al. 2013a |
| **NEARCTIC (sum)** | **296** | **4** | **0.99\*\*\*** |  |
| USA: California | 16 | 0 | 1.00 | Ferrer-Suay et al. 2014a |
| USA: Colorado | 6 | 1 | 0.85 | Ferrer-Suay et al. 2014a |
| USA: Iowa | 2 | 2 | 0.50 | Ferrer-Suay et al. 2014a |
| USA: Georgia | 3 | 0 | 1.00 | Ferrer-Suay et al. 2014a |
| USA: Maryland | 2 | 0 | 1.00 | Ferrer-Suay et al. 2014a |
| USA: Minnesota | 263 | 1 | 0.99*** | Casiraghi et al. 2022 |
| USA: Mississippi | 1 | 0 | 1.00 | Ferrer-Suay et al. 2014a |
| USA: Montana | 1 | 0 | 1.00 | Ferrer-Suay et al. 2014a |
| USA: Utah | 1 | 0 | 1.00 | Ferrer-Suay et al. 2014a |
| USA: Washington | 1 | 0 | 1.00 | Ferrer-Suay et al. 2014a |
| **TOTAL** | **951** | **129** | **0.88\*\*\*** |  |

### *Alloxysta brevis* reproduces via symbiont-mediated thelytokous parthenogenesis

Of 23 field-collected, unmated, *Alloxysta brevis* females reared from *Aphelinus certus* mummies, 16 produced 61 offspring, all of which were daughters (3.8 + 0.9 (SEM) daughters per female).

This result indicated that these parasitoids reproduce via thelytoky, so next we used antibiotic treatment to determine if this reproductive mode was mediated by a bacterium. In total, the antibiotic-treated females produced 2 female and 47 male offspring, while the control females produced 33 females and 10 males (Figure 2). This is a highly significant increase in male production by treated females (Two-tailed Fisher’s exact test; P < 0.0001, Odds Ratio = 71.8 w/ 95% C.I. of 14.8 – 705.3). Furthermore, the daughters produced by antibiotic-treated females were only produced during the first 48-hour oviposition period after antibiotic treatment, while the control females produced daughters throughout the 12-day experimental period (Fig. 2).

**Figure 2.**
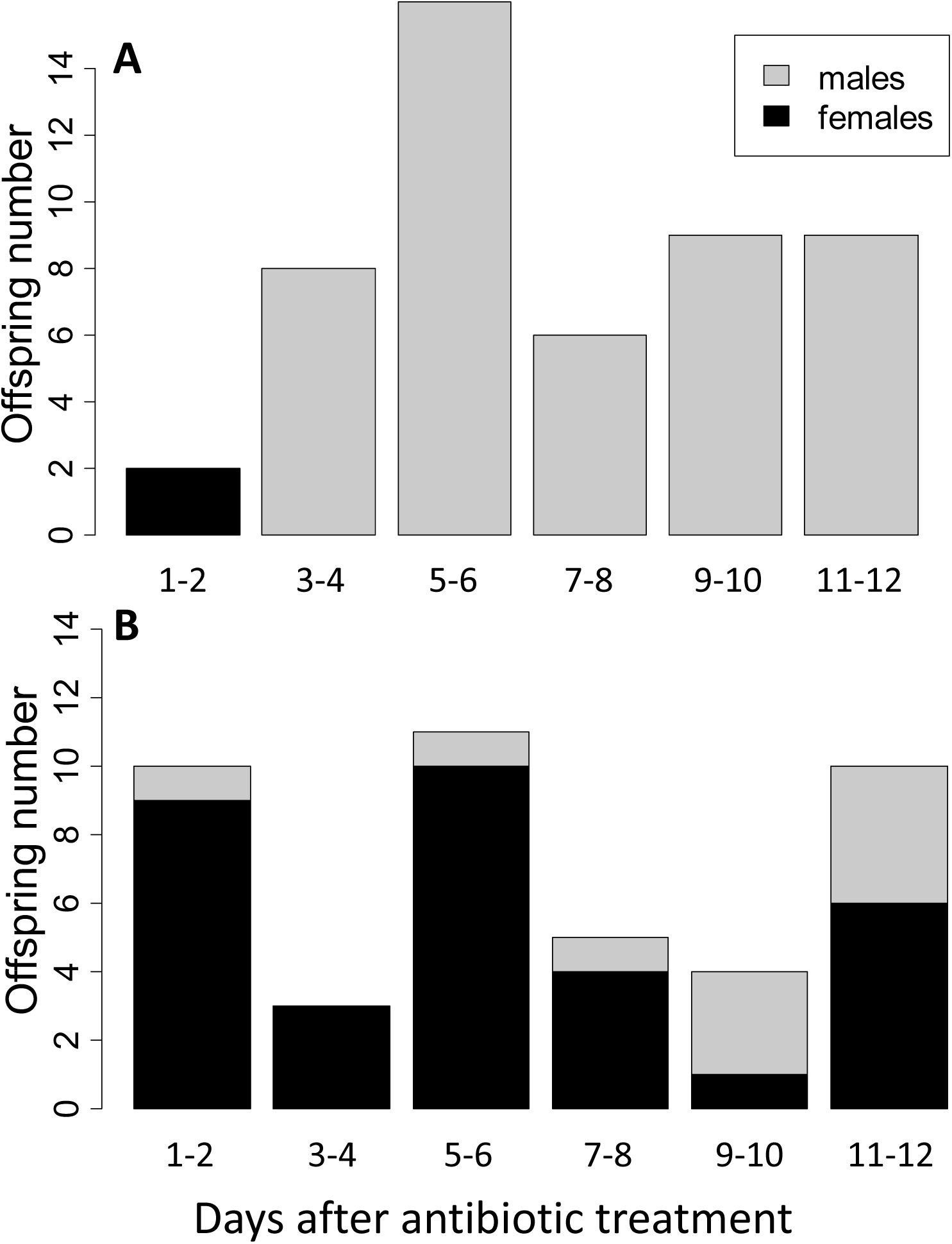
Antibiotic treatment interferes with thelytokous parthenogenesis in *Alloxysta brevis*. **(A)** Numbers of male and female offspring produced by 19 females treated with the antibiotic rifampicin over 12 days after treatment. **(B)** Numbers of male and female offspring produced by 16 control females not treated.

There was a marginally significant trend for a decreasing female-bias over the six time periods in the control group (Two-tailed Fisher’s exact test: *P* = 0.079; Fig. 2b), a pattern that has been noted for other parasitoids that reproduce via symbiont-meditated parthenogenesis (25).

### *Alloxysta brevis* are associated with *Wolbachia* from Supergroup B

*Wolbachia*-specific PCR assays indicated that all *Alloxysta brevis* individuals tested (n=8) were positive for *Wolbachia* DNA (Figure 3). We then used amplicon sequencing of full-length 16S to (1) check for the presence of other symbionts, and (2) determine if there were multiple *Wolbachia* strains present. After removal of environmental contaminants, *Wolbachia* was the only insect-associated microbe in the sequencing data, and assembly of all reads that were assigned to order Rickettsiales resulted in a single high-confidence contig. Given that multiple Rickettsiales (e.g., *Wolbachia* and *Rickettsia*, or perhaps two *Wolbachia* strains) could have been assembled into a single contig, we manually inspected read alignments. Any variants (as compared to the consensus sequence) were (1) present at low and inconsistent frequencies, and (2) did not consistently co-occur in a single read (i.e., there was no support for a second low-frequency haplotype), indicating that there was a single *Wolbachia* strain across all the individuals we screened. Phylogenetic analyses indicated that this *Wolbachia* strain from *Alloxysta brevis*, hereinafter ‘*w*Abre’ is closely related to other *Wolbachia* from ‘Supergroup B’ (Figure 4). The full-length *w*Abre 16S sequence has been deposited in GenBank under accession number PV568413.1.

**Figure 3.**
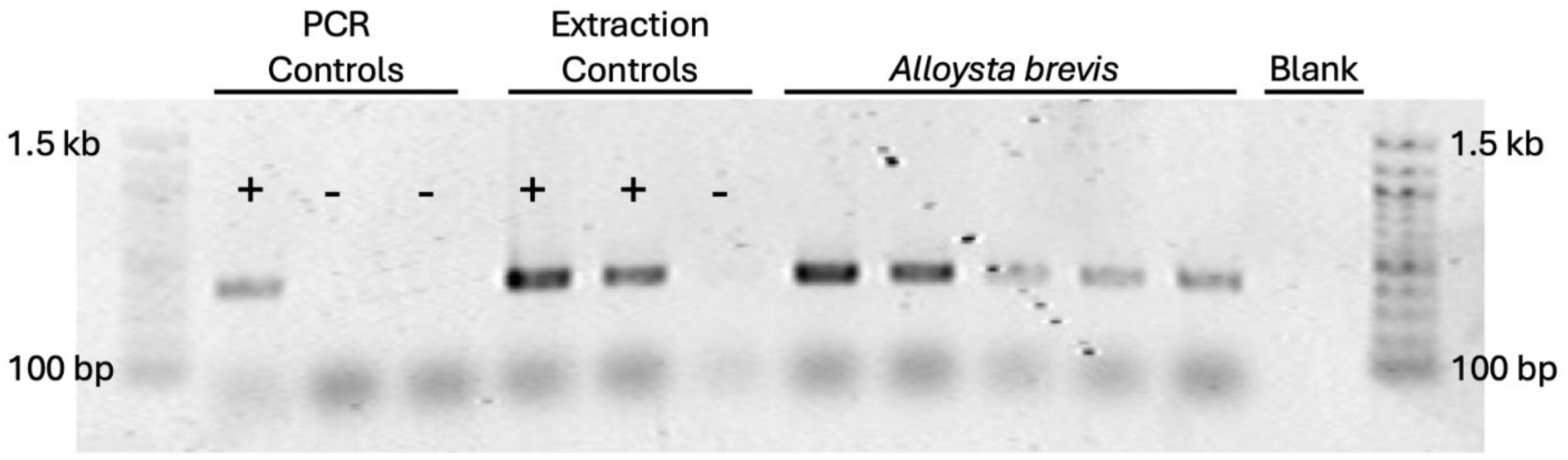
Diagnostic PCR for *Wolbachia*. *Wolbachia*-specific 16S “Wspec” primers (18) were used to test for the presence of *Wolbachia* DNA in individual *Alloxysta brevis* wasps. In total eight individual *Alloxysta brevis* were screened, all positive for *Wolbachia* DNA. The results of five individuals are shown here. PCR controls included (+) a previously tested DNA extraction from *Wolbachia*-infected *Trichogramma pretiosum* (4) and no template (-) reactions. Extraction controls included PCR amplifications of fresh extractions from wasps of the same colony of *Trichogramma pretiosum* (+), and negative extraction controls (-) which included extraction buffers into which the wasp-handling paintbrush was dipped.

**Figure 4.**
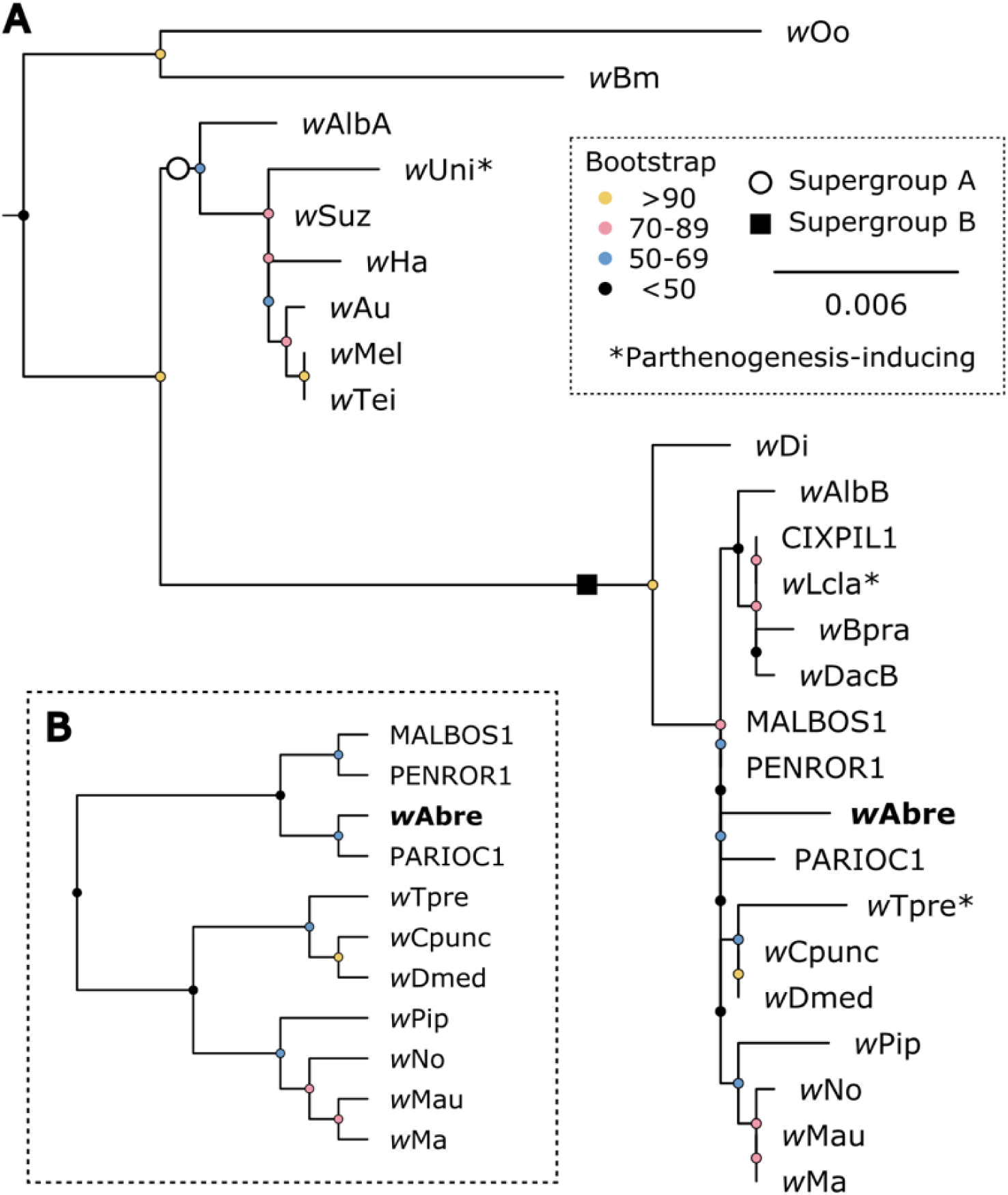
*Wolbachia* strain *w*Abre nests within Supergroup B. **(A)** Phylogenetic reconstruction of *Wolbachia* based on full-length 16S rRNA. **(B)** Monophyletic group within supergroup B containing *w*Abre, displayed as a cladogram to facilitate better interpretation of relationships given the especially short branch lengths in (A).

## DISCUSSION

*Alloxysta brevis* females, collected as immatures in soybean fields in Minnesota, USA, and then isolated in the laboratory with no opportunity to mate, produced only daughters. This result confirms the hypothesis of thelytoky in this population of *Alloxysta brevis*, which had been previously inferred based on an extremely female-biased sex ratio (12). Thelytoky in haplo-diploid species such as the Hymenoptera can be caused by genetic mechanisms or by parthenogenesis-inducing bacterial endosymbionts (1, 2), and we showed that treatment with antibiotics led to the production of male offspring. This demonstrates that bacterial associates are involved in causing thelytoky in this parasitoid species. Further, our molecular analyses identified that the endosymbiotic bacterium *Wolbachia* was associated with *Alloxysta brevis* females. One caveat is that PCR-based assays can result in false positives when *Wolbachia* DNA is integrated into the host genome (26. 27). However, given the results of the antibiotic treatments, and since 16S amplicon sequencing did not recover any other bacterial taxa, we conclude that *Wolbachia* likely mediates parthenogenesis induction in the aphid hyperparasitoid *Alloxysta brevis*. The finding of *Wolbachia*-mediated parthenogenesis induction in *Alloxysta brevis* is not particularly surprising, as at least three members of the family Figitidae exhibit this association (two *Leptopilina* species, and *Gronotoma micromorpha* (Perkins); 2, 26, 28, 29).

Thelytokous parthenogenesis refers specifically to the ability of unmated females to produce daughters and does not exclude the production of males under some conditions. While all of our field-collected *Alloxysta brevis* produced only daughters when held at 24.5°C, the control females in our antibiotic-treatment assay produced appreciable numbers of sons (10 males out of 43 offspring produced). For this second experiment, females in both treatments were held at 26.2°C for 3 days prior to exposure to hosts to improve the effectiveness of the antibiotic rifampicin for the treated parasitoids (see 26, 29, 30). Elevated temperatures are known to ‘cure’ insects of *Wolbachia* endosymbionts, with even temperatures between 25 and 27°C leading to reductions in *Wolbachia* titer (31). We thus hypothesize that elevated temperature was at least partially responsible for male production in the control treatment of this experiment.

Another observation worth noting is that female offspring were produced by treated females over the first two days of the assay. This is not unexpected as the first eggs laid by treated females often do not show signs of endosymbiont elimination in parasitoids (26, 31, 32). This is presumably due to the fact that female parasitoids often eclose as adults with one or more chorionated eggs (33) that would likely be inaccessible to antibiotics ingested by the female.

Furthermore, even if the microbe were eliminated in these mature eggs, the treatment would not eliminate already secreted bacterial effector proteins that drive the developmental changes.

*Alloxysta brevis* exhibits female-biased sex ratios in collections from throughout the world, some of which consist exclusively of females. While our experiments show that *Wolbachia*-mediated thelytoky likely contributes to this global trend, other causes, particularly adaptive patterns of sex allocation by mated females (34) cannot be ruled out. Previous laboratory work with *Alloxysta brevis* using individuals collected in Germany used both females and males, with no mention of thelytoky or female-biased sex ratios (35, 36). This suggests that arrhenotokous populations of *Alloxysta brevis* occur without the intervention of antibiotic treatment. Thus, while we have demonstrated thelytoky in a population of *Alloxysta brevis* collected in Minnesota, USA, the broader literature on this species suggests a scenario in which both thelytokous and arrhenotokous populations exist. This is not unexpected, as it has been seen in other *Wolbachia*-parasitoid interactions such as *Trichogramma* species in California (37, 38). In the Figitidae, *Leptopilina clavipes* (Hartig) includes thelytokous populations in the Netherlands and an arrhenotokous populations in Spain (29). In this case, Dutch parasitoids tested positive for *Wolbachia* while Spanish parasitoids did not. We hypothesize a similar scenario for *Alloxysta brevis*, with the presence of *Wolbachia* (and therefore thelytokous populations) facilitated by conditions that are permissive of *Wolbachia*, or relatively recent local acquisition of *Wolbachia* in some populations.

Since *Alloxysta brevis* is a hyperparasitoid of aphids, its association with parthenogenesis-inducing *Wolbachia* may affect its capacity to disrupt biological control of pest aphid species. Female-biased sex ratios *per se* are expected to increase the effectiveness of parasitoids in suppressing host populations (39–41). For hyperparasitoids, this would suggest a trend for increased disruption of biological control for hyperparasitoids as the female-bias increases. However, harboring parthenogenesis-including *Wolbachia* can also result in lower total reproductive output (42, 43), and the ability of *Wolbachia* to alter offspring sex ratio itself can vary with reproductive rates (25). It is thus not clear how the association with parthenogenesis-inducing *Wolbachia* that we have uncovered for *Alloxysta brevis* should be expected to influence aphid biological control. Further studies comparing populations of this parasitoid that do or do not harbor *Wolbachia* under varying conditions of host availability could shed light on this question. Given the widespread geographic distribution of this parasitoid species, and its broad host range (9, 11), such studies could be of broad relevance to aphid biological control.

## ACKNOWLEDGEMENTS

We thank Nicholas Padowski, Joe Kaser, Kelton Welch and James Miksanek for sampling, Sarah Wood for experimental assistance, and Ulrike Munderloh for supplying the rifampicin. Research reported in this publication was supported by the University of Minnesota Rapid Agricultural Response Fund, the University of Minnesota Agricultural Experiment Station, the Minnesota Invasive Terrestrial Plant and Pest Center, the Minnesota Soybean Research and Promotion Council (all to G.E.H.), and the National Institute of General Medical Sciences of the National Institutes of Health under award number R35GM150991 to A.R.I.L. M.F.-S. was supported by a postdoctoral grant from the Spanish Ministry of Economy and Competitiveness under the contract FJCI201421120 and the project GE 2023 from the Council of Innovation, Universities, Science and Digital Society (CIGE/2022/158).

